# Neuroimaging Measures of Iron and Gliosis Explain Memory Performance in Aging

**DOI:** 10.1101/2021.05.19.444857

**Authors:** Anu Venkatesh, Ana M. Daugherty, Ilana J. Bennett

**Author notes:** Address correspondence to: Anu Venkatesh, Department of Neuroscience University of California, Riverside 900 University Avenue, Riverside CA, 92521-0426.

## Abstract

Evidence from animal and histological studies have indicated that accumulation of iron in the brain results in reactive gliosis that contributes to cognitive deficits. The current study extends these findings to human cognitive aging and suggests that magnetic resonance imaging (MRI) techniques like quantitative relaxometry can be used to study iron and its effects *in vivo*. The effects of iron on microstructure and memory performance were examined using a combination of quantitative relaxometry and multi-compartment diffusion imaging in 35 young (21.06 ± 2.18 years) and 28 older (72.58 ± 6.47 years) adults, who also completed a memory task. Replicating past work, results revealed age-related increases in iron content (R_2_ *) and diffusion, and decreases in memory performance. Independent of age group, iron content was significantly related to restricted (intracellular) diffusion in regions with low-moderate iron (hippocampus, caudate) and to all diffusion metrics in regions with moderate-high iron (putamen, globus pallidus). This pattern is consistent with different stages of iron-related gliosis, ranging from astrogliosis that may influence intracellular diffusion to microglial proliferation and increased vascular permeability that may influence all sources of diffusion. Further, hippocampal restricted diffusion was significantly related to memory performance, with a third of this effect related to iron content; consistent with the hypothesis that higher iron-related astrogliosis in the hippocampus is associated with poorer memory performance. These results demonstrate the sensitivity of MRI to iron-related gliosis and extends our understanding of its impact on cognition by showing that this relationship also explains individual differences in memory performance.

## Introduction

While the neurobiological basis of neurodegeneration and memory decline are multifaceted, iron accumulation and gliosis within gray matter are recognized here as two important contributors. Rather than being independent processes, however, evidence suggests that agerelated accumulation of intracellular unbound, non-heme iron (Hallgren & Sourander, 1958; Mackenzie, Iwasaki, & Tsuji, 2008; Zecca, Youdim, Riederer, Connor, & Crichton, 2004) can promote activation and proliferation of glia (gliosis; Beach, Walker, & McGeer, 1989). For example, *in vitro* (Macco et al., 2013; Pelizzoni, Zacchetti, Campanella, Grohovaz, & Codazzi, 2013) and *in vivo* (Thomsen et al., 2015; You et al., 2017) studies in animal models have directly linked iron-related inflammation to gliosis and subsequent cognitive decline, specifically, memory decline (Schröder, Figueiredo, & De Lima, 2013b; M. Weber et al., 2015). The recently proposed Free-Radical-Induced Energetic and Neural Decline in Senescence (FRIENDS; Raz & Daugherty, 2018) model extends these findings to human cognitive aging and suggests that magnetic resonance imaging (MRI) methods may be sensitive to markers of iron-related gliosis *in vivo* for the study of cognitive aging. The current study applies this model by characterizing the relationships between iron content, gray matter gliosis and memory performance using a combination of quantitative relaxometry and diffusion MRI in young and older adults who also completed a recall memory task.

Although non-heme iron is essential to neurons and glia for key metabolic functions (e.g. adenosine triphosphate production, neurotransmitter synthesis; Zecca et al., 2004), chronic iron related oxidative damage can overwhelm endogenous antioxidant defenses (e.g. glutathione; Vilhardt, Haslund-Vinding, Jaquet, & McBean, 2017) and result in reactive gliosis (Freitas, Ferreira, Trevenzoli, Oliveira, & Reis, 2017; Zecca et al., 2004). This can occur when large concentrations of intracellular iron outside binding complexes (e.g. ferritin; Connor, Menzies, Martin, & Mufson, 1990) produce reactive oxygen species (Macco et al., 2013; Mills et al., 2010), that directly stimulate gliosis (Burda & Sofroniew, 2014). Iron-related oxidative damage and subsequent gliosis drive the cumulative and progressive cognitive declines that are typical in human aging (Raz & Daugherty, 2018). Applying this FRIENDS model, the current study will use individual and age-group differences in gray matter and performance to characterize the nature of the relationship between iron content and gliosis and their joint contributions to memory performance, which has not yet been assessed in humans *in vivo*.

A well-established MRI approach for measuring iron content is R_2_ * relaxometry (Langkammer et al., 2010). This approach has been used in humans to demonstrate age-related accumulation of iron in the basal ganglia and hippocampus (Daugherty, Haacke, & Raz, 2015; Ghadery et al., 2015), consistent with human histological studies (Bartzokis et al., 2007; Zecca et al., 2004). Within the basal ganglia, the globus pallidus has the highest iron concentration across the adult lifespan, whereas the putamen and caudate have a moderate concentration in young adulthood and continue to accumulate iron into old age. This contrasts with the hippocampus, which has less iron concentration in young adulthood and modest accumulation with age (Ghadery et al., 2015). These regional and age group differences in iron content may both affect the relationships between iron and gliosis across the basal ganglia and hippocampus.

Gliosis can have several phenotypes within gray matter, including astrocyte swelling (Singh, Trivedi, Devi, Tripathi, & Khushu, 2016), microglia proliferation (Yi et al., 2019) and increased blood-brain permeability (Simon & Iliff, 2016). The sensitivity of diffusion imaging to these phenotypes has been validated in animal models of age and acute injury (Badaut et al., 2011; Budde, Janes, Gold, Turtzo, & Frank, 2011; Debacker, Djemai, Ciobanu, Tsurugizawa, & Bihan, 2020; Singh et al., 2016; R. A. Weber et al., 2017; Yi et al., 2019; Zhuo et al., 2012) and *in vitro* human (Grussu et al., 2017) studies using a combination of diffusion imaging and histology. Since diffusion imaging is sensitive to different phenotypes of gliosis, it may be used to characterize how different phenotypes and stages (as outlined in Burda & Sofroniew, 2014, Sofroniew, 2015) vary by gray matter region, with the expectation that higher iron would be associated with more pronounced gliosis.

Whereas most of the previous studies have used traditional single-tensor diffusion imaging to investigate gray matter gliosis, the current study will use a multicompartment diffusion approach (Neurite Orientation Dispersion and Density Imaging, NODDI; Zhang, Schneider, Wheeler-Kingshott, & Alexander, 2012). NODDI models diffusion as three separate compartments including restricted (e.g., intracellular), hindered (e.g., extracellular), and free (e.g., cerebral spinal fluid, CSF) diffusion (Zhang et al., 2012, Fukutomi et al., 2018; Kaden, Kelm, Carson, Does, & Alexander, 2016; Rae et al., 2017), which may lead this approach to be more sensitive to gliosis and its different stages across gray matter regions.. When viewed from the perspective of iron-related gliosis, correlations between R_2_ * and NODDI measures across gray matter regions can demonstrate the sensitivity of MRI techniques to regional differences in gliosis staging. For example, in regions with less iron (e.g. hippocampus), early stages of gliosis, including astrocyte swelling (Norenberg, 1994), may be seen as increases in restricted diffusion. In contrast, regions with more iron (e.g. globus pallidus) may also display gliosis associated with microglia proliferation (Yi et al., 2019) and dysregulation of the blood-brain barrier (Andersen, Johnsen, & Moos, 2014), which can increase hindered and free diffusion. Alternatively, regions with the largest age group differences in iron content (e.g., putamen, caudate) may display the most pronounced gliosis compared regions with smaller age group differences in iron (e.g., hippocampus). In either scenario, the hippocampus is likely to have relatively low levels of gliosis that may nonetheless impact cognition in young and older adults given the critical role of this region in memory performance (Lister & Barnes, 2009).

The FRIENDS model of cognitive aging predicts a specific, but as yet untested, combined effect of iron-related gliosis on cognition. Previous studies have separately demonstrated that hippocampal iron content (Ghadery et al., 2015; Rodrigue, Daugherty, Haacke, & Raz, 2013; Schröder et al., 2013a) and microstructure (Carlesimo, Cherubini, Caltagirone, & Spalletta, 2010; Den Heijer et al., 2012; Radhakrishnan, Stark, & Stark, 2020) relate to recall memory performance. Here, we aim to assess the combined influence of hippocampal iron and diffusion to differences in memory performance using a commonality analysis between R_2_ * and NODDI measures.

Building on previous animal research and the FRIENDS model of cognitive aging, the current study aimed to characterize relationships among iron (R_2_*), microstructure (hindered, restricted, free diffusion) and memory performance (RAVLT delayed) in younger and older adults using a multimodal MRI approach. The primary objectives were to: (1) replicate regional and age group differences in iron content, microstructure and memory performance; (2) examine relationships between iron and microstructure in light of the regional and age group differences in iron, and (3) test functional relevance of the iron-microstructure relationship in the hippocampus by examining their contribution to memory performance. Results are expected to show that higher iron concentration (regional difference) and accumulation (age group difference) relates to higher diffusion across the hippocampus and basal ganglia nuclei, with hippocampal iron and microstructure explaining memory performance. Consistent with the FRIENDS model, these results would provide support to the notion that human MRI data can be interpreted using mechanistic hypotheses from the animal research to ultimately better understand cognitive aging.

## Materials and Methods

### Participants

Young and older adults were recruited from the University of California, Riverside (UCR) and surrounding neighborhoods. Prior to enrollment, participants were screened over the phone for neurological conditions (e.g., depression, stroke), scanner related contraindications (e.g., claustrophobia, pregnancy), and general cognition using non-visual portions of the Montreal Cognitive Assessment (MoCA; Nasreddine et al., 2005; Pendlebury et al., 2017). After completing remaining portions of the MoCA in person, all participants exhibited normal cognition with scores > 23 (27.3 ± 1.63). One young and two older participants were excluded due to excessive motion in R_2_* maps. The final sample included 35 young (21.06 ± 2.18 years old, 24 female) and 28 older adults (72.58 ± 6.47 years old, 13 female).

All individuals provided informed consent prior to participation in this study. The UCR Institutional Review Board approved the experimental procedures and participants were compensated for their time.

### Episodic Memory Test

The Rey Auditory Verbal Learning Test (RAVLT; Rey, 1941) was administered to assess delayed free recall, measured as the number of items (out of 15) correctly recalled 30 minutes after completing five immediate free recall trials of the same word list and one immediate free recall trial of a second word list.

### MRI Scanning Protocol

Imaging data were acquired using a 3T Siemens Prisma MRI (Siemens Healthineers, Malvern, PA) scanner fitted with a 32-channel receive-only head coil at the UCR Center for Advanced Neuroimaging.

A high-resolution magnetization-prepared rapid gradient-echo (MP-RAGE) image was acquired with the following parameters: echo time (TE)/repetition time (TR) = 2.72/2400 ms, 208 axial slices, voxel size = 0.8×0.8×0.8 mm, and GRAPPA acceleration factor = 2.

Two diffusion-weighted echo-planar imaging (EPI) sequences were acquired with phase-encoding directions of opposite polarity for correction of susceptibility distortions (Andersson, Skare, & Ashburner, 2003), each with the following parameters: TE/TR = 102/3500 ms, FOV = 212×182 mm, matrix size of 128×110, voxel size = 1.7×1.7×1.7 mm, 64 axial slices, and multiband acceleration factor = 4. For each acquisition, bipolar diffusion encoding gradients (b = 1500 and 3000 s/mm^2^) were applied in 64 orthogonal directions, with six images having no diffusion weighting (b = 0; 12 total).

Multi-echo data derived from a 12-echo 3D gradient recalled echo (GRE) sequence were acquired with the following parameters: TE/ΔTE/TR = 4/3/40 ms, FOV = 192×224 mm, matrix size = 192×224×96, slice thickness = 1.7 mm, and GRAPPA acceleration factor = 2. Magnitude and phase images were saved for later calculation of R_2_* values.

### Region of Interest Segmentations

Bilateral hippocampus, caudate, putamen, and globus pallidus were automatically segmented on each participant’s MP-RAGE image using FMRIB Software Library’s (FSL; Jenkinson, Beckmann, Behrens, Woolrich, & Smith, 2012) Integrated Registration and Segmentation Tool (FIRST; Patenaude, Smith, Kennedy, & Jenkinson, 2011), with the flag for three-stage affine registration for hippocampus (as in Venkatesh et al., 2020). After visual inspection of each region of interest (ROI), caudate segmentations that underestimated the structure were corrected using a flag to increase the number of modes of variation for fitting from the default (40) to the maximum (336; n = 4 young) and those that were misaligned were corrected using a linear registration between the MP-RAGE and standard brain (Montreal Neurological Institute; MNI) instead of the default subcortical mask (n = 1 young). No corrections were needed for the hippocampus, putamen or globus pallidus segmentations.

### Iron Image Processing

For each participant, iron data were pre-processed using the procedure outlined in Langley et al. (2019). Briefly, R_2_* values were estimated using a custom script in MATLAB which fit a monoexponential model, (*Si* = *S0*exp [−*R*_2_*TE], where *Si* indicates the signal of a voxel at the *i*th echo time and *S0* indicates a fitting constant) to the GRE images.

FSL’s FMRIB Linear Image Registration Tool (FLIRT) was used to align the resulting R_2_* map to the MPRAGE image via the magnitude image from the first echo, using a rigid body transformation (six degrees of freedom, DOF). An affine transformation (12 DOF) with nearest neighbor interpolation was used to align the FIRST segmented ROIs into iron space using FLIRT and the matrix file from the previous step. Each bilateral iron space-aligned ROI mask was then multiplied by the voxel-wise R_2_* map before taking the average across voxels and mean R_2_* was extracted for each participant,

For each bilateral ROI, mean R_2_* (Iron_raw_) was adjusted for ROI volume using the normalization method from Jack et al. (1989). The FIRST-segmented ROI volumes (Volume_indiv_) were used to calculate adjusted R_2_* (Iron_norm_) separately for each participant using the following equation: Iron_norm_ = Iron_raw_ − β (Volume_indiv_ − Volume_mean_). Mean volume (Volume_mean_) and slope (β) were calculated within the young adults. Volume-adjusted R_2_* values were used for all analyses.

### Diffusion Data Processing

For each participant, diffusion data were pre-processed using FSL, except that a binary brain mask was created using Analysis of Functional Neuro Images (AFNI; Cox 1996). After generating a field map using Topup, Eddy was used to correct for distortions due to motion, eddy-currents, and susceptibility (Andersson, Skare, & Ashburner, 2003; Andersson & Sotiropoulos, 2016).

The NODDI MATLAB toolbox was then used to estimate voxel-wise measures of restricted (also known as intracellular volume fraction, ICVF), hindered (also known as orientation dispersion index, ODI) and free (also known as isotropic fraction, fISO) diffusion (http://mig.cs.ucl.ac.uk/index.php; Zhang, Schneider, Wheeler-Kingshott, & Alexander, 2012). To more accurately model diffusion within gray matter, the intrinsic diffusivity assumption, used to estimate restricted and hindered diffusion, was set to 1.1 × 10^−3^ mm^2^/s (Guerrero et al. 2019, Fukutomi et al., 2019, 2018).

For each participant, diffusion metrics were extracted separately for each FIRST segmented ROI. A rigid body transformation was used to align the FIRST segmented ROIs to diffusion space using FLIRT. For free diffusion, a bilateral diffusion space-aligned ROI mask was multiplied by the voxel-wise free diffusion image before taking the average across voxels. To limit hindered and restricted diffusion metrics to voxels with sufficient tissue content, an inclusion mask was created by thresholding the free diffusion image to voxels with high tissue content (free diffusion < 90%). The inclusion mask was then multiplied by each bilateral diffusion space-aligned ROI mask and then by the corresponding voxel-wise diffusion image before taking the average across voxels and hemispheres.

### Statistical Analyses

Repeated measures ANOVAs (main effects; M ± SD, interactions; M ± SEM), *t*-tests and regression analyses were conducted using SPSS (Version 24.0; IBM, Armonk, NY, USA). For all analyses, the significance threshold was set to *p* < 0.05, unless otherwise noted.

To test for neural correlates of the memory measure, it is equally important to consider the correlated effect of iron content and gliosis as well as their unique effects. To determine the shared effect, a commonality analysis was performed (Lindenberger, von Oertzen, Ghisletta, & Hertzog, 2011). The commonality analysis uses a series of linear regressions to calculate the shared and unique effects of each predictor (iron content and diffusion) and estimates the shared variance of the predictors as a proportion of the total variance explained in memory performance (shared over simple effect; SOS). Large values would indicate high commonality between predictors, which is consistent with the iron-gliosis model reviewed.

## Results

### Iron Content

An Age Group (young, older) × Region (hippocampus, caudate, putamen, globus pallidus) repeated measures ANOVA was conducted for R_2_* (Figure 1). There was a significant effect of Region, *F*(3, 183) = 492.36, *p* < 0.001, with the highest iron content in the globus pallidus (33.46 ± 4.72), followed by the putamen (24.27 ± 4.84), caudate (20.97± 2.57), and hippocampus (16.40 ± 1.49). Post hoc pairwise comparisons revealed that iron content was significantly different between all regions, *p*s < 0.001.

**Figure 1.**
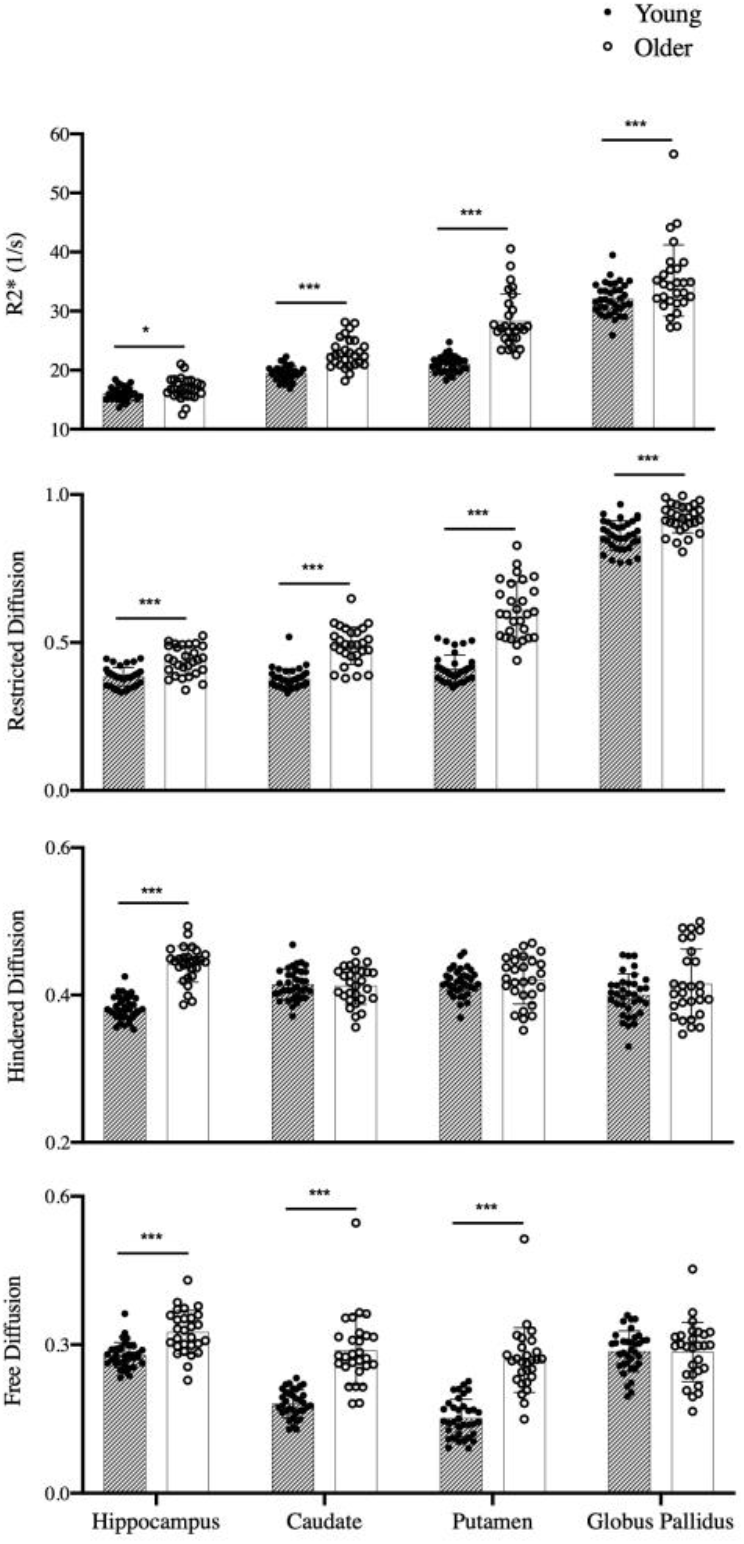
Iron content (R_2_*) and microstructure (restricted, hindered, free diffusion) are shown separately for young (black circles, stripe bar) and older (open circles, open bar) adults in each region of interest. *** *p* < 0.001, * = *p* < 0.05

There were also significant effects of Age Group, *F*(1, 61) = 51.47, *p* < 0.001, and Age Group × Region, *F*(3, 183) = 16.77, *p* < 0.001. Overall, iron content was higher in older adults (25.81 ± 0.38) compared to young (22.15 ± 0.34). Post hoc 2 Age Group × 2 Region comparisons revealed that the age group difference was significantly larger in the putamen (7.29 ± 0.81) compared to the caudate (3.46 ± 0.49), *F*(1, 61) = 40.10, *p* < 0.001, globus pallidum (3.06 ± 1.14), *F*(1, 61) = 13.67, *p* < 0.001, and hippocampus (0.83 ± 0.37), *F*(1, 61) = 49.27, *p* < 0.001; and in the caudate compared to the hippocampus, *F*(1, 61) = 49.27, *p* < 0.001. The age-related differences in globus pallidus were statistically equivalent to that in the caudate (*p* > 0.20) and hippocampus (*p* > 0.05).

### Microstructure

Age Group (young, older) × Region (hippocampus, caudate, putamen, globus pallidus) repeated measures ANOVAs were conducted separately for each diffusion metric. In the event of significant interactions, post-hoc Age Group × Region ANOVAs for each pair of regions was conducted.

#### Restricted Diffusion

There was a significant effect of Region, *F*(3, 183) = 1851.3, *p* < 0.001, with the highest restricted diffusion in the globus pallidus (0.89 ± 0.06), followed by the putamen (0.50 ± 0.12), caudate (0.43 ± 0.07), and hippocampus (0.41 ± 0.05). Post hoc pairwise comparisons revealed that restricted diffusion was significantly different between all regions, *p*s < 0.008.

There were significant effects of Age Group, *F*(1, 61) = 102.84, *p* < 0.001, and Age Group × Region, *F*(3, 183) = 41.05, *p* < 0.001. Overall, restricted diffusion was higher in older adults (0.61 ± 0.01) compared to young (0.51 ± 0.01). Post hoc 2 Age Group × 2 Region comparisons revealed that the age group difference was significantly larger in the putamen (0.19 ± 0.02) compared to the caudate (0.11 ± 0.01), *F*(1, 61) = 46.97, *p* < 0.001, globus pallidus (0.06 ± 0.01), *F*(1, 61) = 78.73, *p* < 0.001, and hippocampus (0.05 ± 0.01), *F*(1, 61) = 65.11, *p* < 0.001; in the caudate compared to the globus pallidus (0.05 ± 0.01), *F*(1, 61) =14.95, *p* < 0.001, and hippocampus, *F*(1, 61) = 20.76, *p* < 0.01; and in the globus pallidus compared to the hippocampus, *F*(1, 61) =0.09, *p* < 0.001.

#### Hindered Diffusion

The effect of Region was not significant, *p* > 0.08, but there were significant effects of Age Group, *F*(1, 61) = 16.10, *p* < 0.001, and Age Group × Region, *F*(3, 183) = 21.63, *p* < 0.001. Overall, hindered diffusion was higher in older adults (0.42 ± 0.01) compared to young (0.40 ± 0.01). Post hoc 2 Age Group × 2 Region comparisons revealed that the age group difference was significantly larger in the hippocampus (0.06 ± 0.01) compared to the caudate, *F*(1, 61) = 74.63, *p* < 0.001, putamen, *F*(1, 61) = 58.46, *p* < 0.001, and globus pallidus, *F*(1, 61) = 19.84, *p* < 0.001. The results did not statistically differ between the remaining regions, *p*s > 0.09.

#### Free Diffusion

There was a significant effect of Region, *F*(3, 183) = 81.37, *p* < 0.001, with the highest free diffusion in the hippocampus (0.30 ± 0.04), followed by globus pallidus (0.29 ± 0.05), caudate (0.23 ± 0.07) and putamen (0.20 ± 0.08). Post hoc pairwise comparisons revealed that free diffusion was significantly different between all regions, *p*s < 0.001, except between globus pallidus and hippocampus, *p* > 0.23.

There were also significant effects of Age Group, *F*(1, 61) = 58.14, *p* < 0.001, and Age Group × Region, *F*(3, 183) = 34.22, *p* < 0.001. Overall, free diffusion was higher in older (0.29 ± 0.01) compared to young (0.22 ± 0.01) adults. Post hoc 2 Age Group × 2 Region comparisons revealed that the age group difference was significantly larger in the hippocampus (0.05 ± 0.02) compared to the globus pallidus (0.01 ± 0.01), *F*(1, 61) = 9.66, *p* < 0.004, caudate (0.11 ± 0.04), *F*(1, 61) = 31.74, *p* < 0.001, and putamen (0.12 ± 0.03), *F*(1, 61) = 61.00, *p* < 0.001; and in the globus pallidus compared to the caudate, *F*(1, 61) = 40.81, *p* < 0.001, and putamen, *F*(1, 61) = 58.92, *p* < 0.001. The results did not statistically differ between the remaining regions, *p*s > 0.30.

### Relation between Iron and Microstructure

Separate linear regressions for each region tested the relationship between iron content (R_2_*) and each diffusion metric, as well as the potential moderating effect of age group (by including Age Group × R_2_* as a predictor). Age group was included as a covariate in all models given the previously described age effects. Significant effects were Bonferroni corrected for three comparisons per diffusion metric (*p* < 0.02; Figure 2).

**Figure 2.**
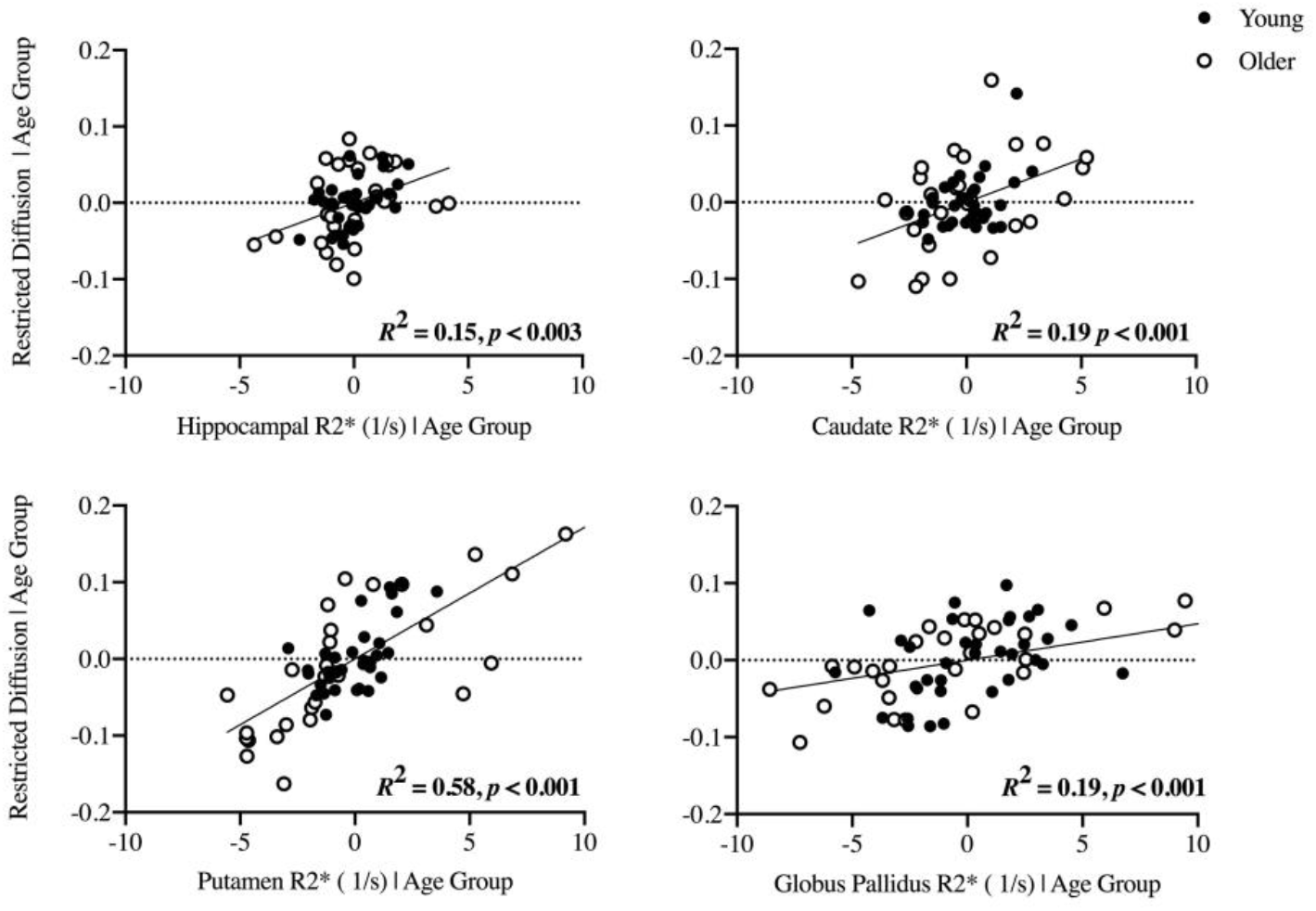
Significant associations between iron content (R_2_*) and microstructure (restricted, hindered, free diffusion) are shown separately for each region after controlling for age group.

For the hippocampus, ß = 0.34, *t*(62) = 3.31, *p* < 0.003, and caudate, ß = 0.39, *t*(62) = 3.75, *p* < 0.001, significant positive relationships were observed between R2* and restricted diffusion, but not hindered or free diffusion, *p*s > 0.03. For the putamen, significant positive relationships were observed between R_2_* and restricted, ß = 0.67, *t*(62) = 8.85, *p* < 0.001, hindered, ß = 0.95, *t*(62) = 6.23, *p* < 0.001, and free, ß = 0.54, *t*(62) = 4.98, *p* < 0.001, diffusion. For the globus pallidus, significant positive relationships were observed between R_2_* and restricted, ß = 0.40, *t*(62) = 3.83, *p* < 0.001, and hindered, ß = 0.52, *t*(62) = 4.52, *p* < 0.001, diffusion, whereas a significant negative relationship was observed between R_2_* and free diffusion, ß = −0.47, *t*(62) = −3.83, *p* < 0.001. There was no evidence of age group moderating these relationships in any region, *p*s > 0.10, indicating that the R_2_*-diffusion relationship was comparable in young and older adults.

### Contributions of Iron and Microstructure to Memory Performance

An independent sample *t*-test assessed age group differences in RAVLT delayed recall, *t*(47) = −4.06, *p* < 0.001, 95% CI [−4.90, −1.66]. As expected, older adults (8.04 ± 3.59) recalled significantly fewer words than young (11.31 ± 2.57).

A commonality analysis quantified the shared variance between hippocampal iron (R_2_*) and microstructure (restricted diffusion) in explaining in memory performance (delayed free recall). These analyses were limited to the hippocampus due to its known role in memory and to the restricted diffusion metric given its previously described relationship to hippocampal iron. Results revealed that 31.5% of the variance in delayed recall performance was explained by restricted diffusion alone, 11.8% by R_2_* alone, and a total of 32.4% when both restricted diffusion and R_2_* were included in the model (see Table 1). From this procedure, of the total variance in RAVLT recall that was explained by diffusion, 34.6% of the effect was shared with hippocampal R_2_*.

**Table 1.**
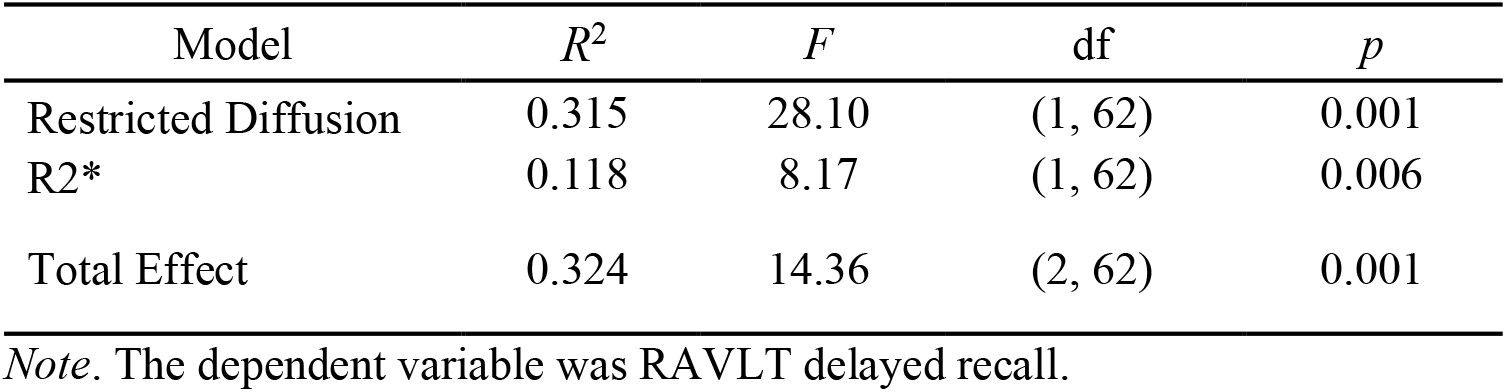
Summary of regression models for the commonality analysis.

| Model | $R^2$ | $F$ | df | $p$ |
| --- | --- | --- | --- | --- |
| Restricted Diffusion | 0.315 | 28.10 | (1, 62) | 0.001 |
| R2* | 0.118 | 8.17 | (1, 62) | 0.006 |
| Total Effect | 0.324 | 14.36 | (2, 62) | 0.001 |

*Note.* The dependent variable was RAVLT delayed recall.

Since the variance in delayed recall performance explained by our metrics of interest may be shared with age, the commonality analysis was repeated after including age group as a covariate (see Figure 3). In this model, restricted diffusion uniquely explained 12.7% of variance in delayed recall, *R*^2^ = 0.13, *p* < 0.001, the unique effect of R_2_* was 4.8%, *R*^2^ = 0.05, *p* = 0.05, and the total effect of both predictors was 13.5%. Therefore, even independent of age group, 31.5% of the total variance in delayed recall that was explained by hippocampal restricted diffusion was shared with hippocampal R_2_*. Taken together, hippocampus microstructure significantly contributes to memory performance independent of age, and approximately 32% of its effect is also related to iron content.

**Figure 3.**
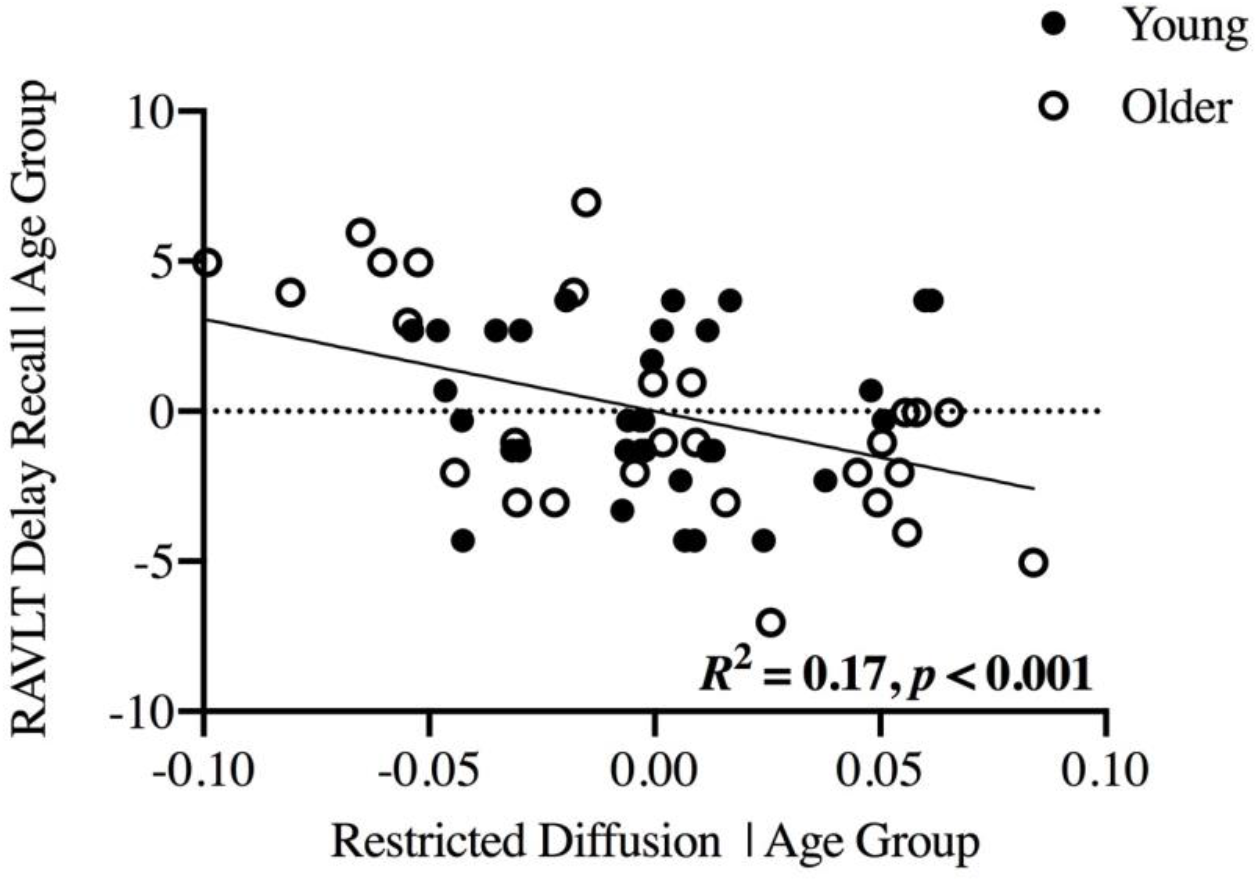
Significant association between hippocampal restricted diffusion and RAVLT delayed recall performance, independent of age group.

## Discussion

The current study tested the relationships between iron, gliosis, and memory in humans using a combination of neuroimaging techniques, consistent with the FRIENDS model of cognitive aging. Our results revealed several major findings. First, we replicated well-known regional and age group differences in iron content (R_2_*), tissue microstructure (NODDI) and memory performance (RAVLT delayed recall). Second, we observed relationships between iron and microstructure that were specific to restricted diffusion in the hippocampus and caudate, whereas they were observed in all diffusion measures in the putamen and globus pallidus, consistent with stages of gliosis as a function of regional iron content. Moreover, these iron-microstructure relationships were not moderated by age group, suggesting that the effect of iron on microstructure may be cumulative and progressive across the adult lifespan. Third, restricted diffusion in the hippocampus related to recall memory performance, with a third of this variance shared with iron. These results demonstrate that MRI is sensitive to iron-related gliosis within gray matter, which contributes to individual differences in memory performance.

As expected, we observed age group differences in delayed recall (Bennett, Huffman, & Stark, 2015), gray matter microstructure (Nazeri et al., 2015; Radhakrishnan et al., 2020; Venkatesh, Stark, Stark, & Bennett, 2020), and iron content (Bartzokis et al., 2007) that replicated previous studies. Of particular importance given our interest in relationships between iron and diffusion measures, we observed the expected patterns of regional and age group differences in iron. That is, iron concentration was highest in the globus pallidus and putamen followed by the caudate and hippocampus, whereas age-related iron accumulation was largest in the putamen and caudate followed by the globus pallidus and hippocampus. This allowed us to examine the extent to which NODDI metrics are sensitive to various stages of iron-related gliosis across the gray matter regions of interest.

We further observed significant relationships between iron content and microstructure that varied across the hippocampus and basal ganglia nuclei. Within the hippocampus and caudate, relationships between R_2_*and NODDI metrics were specific to restricted diffusion. Of note, these regions had low to moderate overall iron concentration in spite of having both small (hippocampus) and large (caudate) are group differences in iron accumulation. We interpret this pattern of results as being consistent with an earlier stage of iron-related gliosis (Norenberg, 1994; Pekny & Nilsson, 2005), in which astrocyte activation and swelling are limited to the intracellular source of diffusion. The positive direction of these effects also supports the notion that higher iron content is accompanied by reactive astrogliosis through oxidative damage, and hence an increase in intracellular sources of diffusion. Proposing astrogliosis as a potential mechanism that influences restricted diffusion extends previous work that has traditionally attributed this diffusion metric to neurite density (Fukutomi et al., 2019; Grussu et al., 2017; Metzler-Baddeley et al., 2019; Radhakrishnan et al., 2020) and provides a parsimonious explanation for previous observations of age-related *increases* in gray matter restricted diffusion seen by our group (Franco et al., 2020; Venkatesh et al., 2020) among others (Metzler-Baddeley et al., 2019; Radhakrishnan et al., 2020).

In contrast, within the putamen and globus pallidus, R_2_* was related to all three diffusion metrics. These regions had moderate to high overall iron concentration, but both large (putamen) and small (globus pallidus) are group differences in iron accumulation. This pattern of results may indicate later stages of iron-related gliosis in which astrogliosis is coupled with microglia proliferation (Yi et al., 2019) and increased vascular permeability (Elahy et al., 2015) that would influence extracellular and free, not just intracellular, sources of diffusion. Recent evidence supports the notion that hindered diffusion is sensitive to infiltrating microglia, as one study demonstated that hindered diffusion significantly varied depending on microglia density in mice (Yi et al., 2019). Whereas R_2_* was only positively related to restricted and hindered diffusion, positive (putamen) and negative (globus pallidus) correlations were seen for free diffusion, which likely reflects low signal to noise ratios in the diffusion signal within the globus pallidus. Taken together, the regional patterns between iron content and microstructure observed here appear to reflect an iron concentration-dependent effect on microstructure. As such, our findings are consistent with and extend the iron-gliosis hypothesis in humans by demonstrating that increased iron accumulation in gray matter is accompanied by a glial response that can be detected initially with intracellular (restricted) and then extracellular (hindered, free) diffusion metrics. Further, by finding that the iron-microstructure relationships were comparable between young and older adults across the hippocampus and all basal ganglia nuclei, our results suggest that iron-related gliosis is cumulative and progressive across the lifespan.

Finally, we demonstrated the shared consequence of hippocampal iron-related gliosis on recall memory performance, providing functional relevance to the current findings. Greater hippocampal restricted diffusion explained 31.5% of the variance in memory performance, 34.6% of this effect was shared with hippocampal R_2_*. Consistent with our interpretation of a cumulative effect of iron across the lifespan, approximately 31.5% of shared variance between microstructure and iron estimates remained after statistically controlling for age. These findings extend at least one previous study that observed that higher restricted diffusion related to poorer memory performance in younger and older adults (Radhakrishnan et al., 2020) by revealing the extent to which this diffusion-memory relationship is shared with iron. More importantly, these behavioral results provide an important piece of evidence in support of the iron-gliosis hypothesis and FRIENDS model by demonstrating the sensitivity of MRI to iron-related hippocampal astrogliosis as a correlate to memory performance.

In conclusion, the current study revealed key pieces of evidence in support of the iron-gliosis hypothesis in humans. We found significant relationships between iron content and tissue microstructure that systematically varied across subcortical regions (but not age groups) in an iron concentration-dependent manner. This has functional consequences as iron content and tissue microstructure together contribute to recall memory performance, independent of age. This study represents an important validation and extension of both the animal literature that gave rise to the iron-gliosis hypothesis and the FRIENDS model by demonstrating that MRI is sensitive to individual differences in iron and gliosis, and that their combined effect explains memory performance.

